# The *Helicobacter pylori* Methylome is Acid-Responsive due to Regulation by the Two-Component System ArsRS and the Type I DNA Methyltransferase HsdM1 (HP0463)

**DOI:** 10.1101/2023.09.22.558991

**Authors:** Elise H. Zimmerman, Erin L. Ramsey, Katherine E. Hunter, Sarah M. Villadelgado, Celeste M. Phillips, Ryan T. Shipman, Mark H. Forsyth

## Abstract

In addition to its role in genome protection, DNA methylation can regulate gene expression. In this study, we characterized the impact of acidity, phase variation, and the ArsRS TCS on the expression of the Type I m6A DNA methyltransferase HsdM1 (HP0463) of *Helicobacter pylori* 26695 and their subsequent effects on the methylome. Transcription of *hsdM*1 increases at least 4-fold in the absence of the sensory histidine kinase ArsS, the major acid-sensing protein of *H. pylori*. *hsdM*1 exists in the phase-variable operon *hsdR*1-*hsdM*1. Phase-locking *hsdR*1 (HP0464), the restriction endonuclease gene, has significant impacts on the transcription of *hsdM*1. To determine the impacts of methyltransferase transcription patterns on the methylome, we conducted methylome sequencing on samples cultured at pH 7 or pH 5. We found differentially methylated motifs between these growth conditions, and that deletions of *arsS* and/or *hsdM*1 interfere with the epigenetic acid response. Deletion of *arsS* leads to altered activity of HsdM1 and multiple other methyltransferases under both pH conditions indicating that the ArsRS TCS, in addition to direct effects on regulon transcription during acid acclimation, may also indirectly impact gene expression via regulation of the methylome. We determined the target motif of HsdM1 (HP0463) to be the complementary bipartite sequence pair 5’-(HH)TCA^m6A^VN_6_TGY-3’ and 3’-AGTN_6_GA^m6A^CA-5’. The Type II m5C DNA methyltransferase M.HpyAVIII (HP1121) is regulated by ArsS and HsdM1. This complex regulation of DNA methyltransferases, and thus differential methylation patterns, may have implications for the decades-long persistent infection by *H. pylori*.

**Importance:** This study expands the possibilities for complex, epigenomic regulation in *Helicobacter pylori*. We demonstrate that the *H. pylori* methylome is plastic and acid-sensitive via the two-component system ArsRS and the DNA methyltransferase HsdM1. The control of a methyltransferase by ArsRS may allow for a layered response to changing acidity. Likely, an early response whereby ArsR∼P affects regulon expression, including the methyltransferase *hsdM*1. Then, a somewhat later effect as the altered methylome, due to altered HsdM1 expression, subsequently alters the expression of other genes involved in acclimation. The intermediate methylation of certain motifs supports the hypothesis that methyltransferases play a regulatory role. Untangling this additional web of regulation could play a key role in understanding *H. pylori* colonization and persistence.

## Introduction

*Helicobacter pylori* is a gram-negative spiral-shaped bacterium that colonizes the gastric epithelium and can cause gastritis, gastric and duodenal ulcers, and gastric adenocarcinoma^1,2^. *H. pylori* is one of the most common bacterial infections in the world, infecting about half of the world’s population^2^. Infection generally occurs in childhood and persists for the host’s lifetime if untreated^3^. Infection can occur via oral-oral transmission, fecal-oral transmission, or can be food-borne or water-borne^4^. While most cases are asymptomatic, 10-15% of infections lead to peptic ulcer disease or gastric cancer^5^. *H. pylori* is the leading cause of gastric adenocarcinoma, which is the second leading cause of cancer-related deaths worldwide^3,5^.

As the pH of the human stomach drastically fluctuates throughout the day and lifetime of the host, acid acclimation strategies are vital for *H. pylori* long-term survival and colonization^6,7,8^. The main acid-sensing and response system in *H. pylori* is the two-component system (TCS) ArsRS which exhibits widespread gene regulation^9,10^. Multiple studies have investigated the global changes in gene expression in response to acid conditions and agree that ArsRS has a regulon of at least 100 genes in multiple *H. pylori* strains^7,8,9,11,12^.

Restriction-modification (R-M) systems are best known for their role as primitive bacterial immune systems in which DNA methylation is a mechanism to label the genome as self^13^. There is a growing body of research demonstrating that R-M systems have functions beyond genome protection, particularly in pathogenesis^1,14,15,16,17,18^. The traditional function of degrading foreign DNA may not provide sufficient explanation for the high level of specificity in sequence recognition, the diversity in kind and number of R-M systems in the bacterial domain, or the independent evolution of restriction endonucleases and methyltransferases in relation to each other^14^.

There are three classes of restriction-modification (R-M) systems in *H. pylori*: Types I, II, and III^19^. They differ in subunit composition of their enzymes, motif recognition, and recombination ability^13^. This study focuses on a Type I DNA methyltransferase in *Helicobacter pylori* strain 26695, HsdM1 (HP0463). Type I R-M systems are complexes composed of three subunits: a DNA methyltransferase (MTase), a restriction endonuclease (REase), and a specificity subunit (S)^20,21^. Type I R-M systems control sequence recognition via two target recognition domains, TRD1 and TRD2, within the S subunit that together allow the protein to recognize a bipartite target sequence^22^. The TRDs are flanked by repeat sequences, the length of which determines the distance between the target sequences^22^. Type II R-M systems are the most abundant and well-characterized class, thus are used as the control in this study. Type II R-M systems are composed of separate REase and DNA MTase subunits which exhibit identical DNA binding specificity^23^. Type II systems bind short, generally palindromic, motifs^13^.

There are three types of bacterial DNA methylation: N6-methyladenosine (m6A), N4-methylcytosine (m4C), and N5-methylcytosine (m5C)^9^. Adenosine methyltransferases are the most prevalent because m5C methylation is highly susceptible to C to T mutations via deamination^24^. Deamination leads to increased mutation rates and decreased DNA stability^14,24^. Thus, bacteria have evolved the utilize m6A and m4C more frequently to avoid these impacts^25^.

Within *H. pylori’s* relatively small genome of 1.64–1.67 Mb, there are over 20 restriction-modification systems^19, 26^. This makes *H. pylori* an organism of interest for R-M system studies as the average number of MTase genes in a prokaryotic genome is five^27^. Methylome analysis across 541 *H. pylori* genomes found that methylation patterns are directly subject to natural selection and linked phenotypes to specific methylation sites^28^. In a comparative genomics study between *H. pylori* strains 26695 and J99, the two earliest sequenced isolates, nine Type II MTases were conserved across both strains without the presence of cognate restriction enzymes^26^. Our study was conducted using *H. pylori* strain 26695 and focuses primarily on its Type I R-M systems along with two selected Type II systems. *H. pylori* strain 26695 possesses three Type I MTases, all m6A-specific, 16 Type II MTases of all three methylation types, and three Type III MTases, all m6A-specific^19^. As these systems are over-represented in the *H. pylori* genome, the possibilities for complex genome regulation are significant.

DNA methylation has been hypothesized to disrupt simultaneous transcription and translation in bacteria by changing the secondary structure of DNA and potentially physically blocking the binding of the NusG/RNA polymerase and NusG/ribosome complexes^29^. Methylation by Type I R-M systems has been shown to decrease transcription of operons, and evidence shows that methylation at start and stop codons can increase or decrease transcription respectively ^30,15^. Type I R-M systems have been shown to be capable of altering their sequence specificity via homologous recombination and point mutations, as well as methylating promoter sequences of other R-M systems in a hierarchical manner^27^. This further suggests that bacteria use these DNA methyltransferases as a way of finely tuning their gene expression levels, potentially allowing them to react in more nuanced manners to a changing environment^30^.

The principal operon under investigation in the current study is *hsdR*1-*hsdM*1 (locus tags HP0464-0463) within *H. pylori* strain 26695, which consists of genes for the REase and DNA MTase subunits, respectively, of a Type I restriction-modification system^31^. HsdM1 has been demonstrated to be regulated by at least three other DNA MTases in *H. pylori* strains 26695 and P12^27^. In addition, this operon is phase-variable based upon a repetitive nucleotide sequence in *hsdR*1^31^. Across all *H. pylori* strains, restriction-modification genes are phase-variable at a disproportionately high rate^19^. Phase variable R-M systems have been shown to switch activity during the early infection stages^31,32,33^. This strongly suggests that phase-variation has evolved to allow *H. pylori* to respond to the various constantly changing environments of the stomach, likely through transcriptional regulation, which is critical to the long-term persistence characteristic of *H. pylori* infection^31,32,33^. Thus, R-M systems are likely a mechanism through which the bacterium control gene expression which may play a key role in colonization and persistence.

## Results

### *hsdM*1 transcription is acid-responsive

The Type I DNA methyltransferase *hsdM*1 (HP0463) exists in the same operon as its cognate restriction endonuclease gene *hsdR*1 (HP0464) while *hsdS*1b (HP0462), the cognate specificity subunit of this Type I restriction-modification system, has its own promoter^34^ (**Fig. 1**). *hsdM*1 transcription is significantly upregulated in the absence of the *H. pylori* 26695 major acid-sensing protein, the sensory histidine kinase ArsS (**Fig. 2a**), but was unaffected by the absence of either of the other two sensory histidine kinases, CrdS or FlgS (data not shown). These data demonstrate that *hsdM*1 is controlled by the two-component system (TCS) ArsRS.

**Figure 1.**
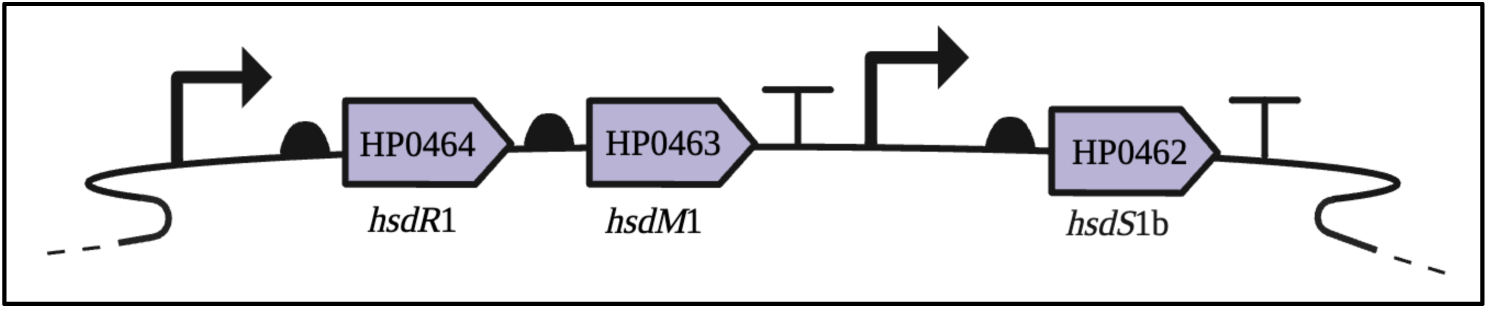
Type I R-M System hsdR1-hsdM1-hsdS1b (HP0464-0462) in the H. pylori 26695 genome. The organization of the restriction-modification system operon HP0464-0463 (hsdM1-hsdR1) and its cognate specificity unit HP0462 (hsdS1b) in the H. pylori 26695 genome. The transcription start sites and presumed terminators are based on Sharma et al., 2010^34^. Figure made in BioRender.

**Figure 2.**
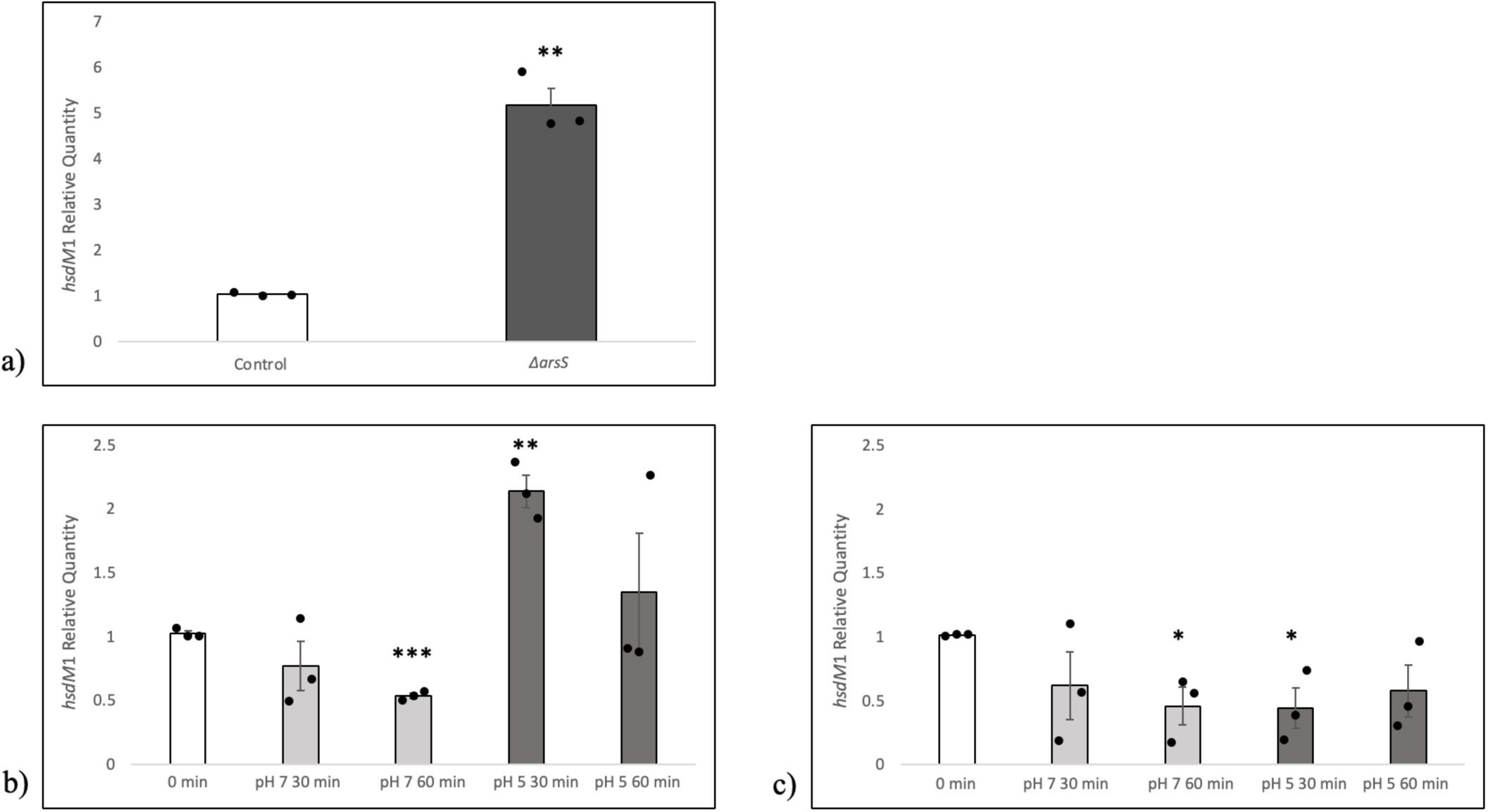
hsdM1 transcription is acid-responsive in an ArsS-dependent manner. **a)** hsdM1 mRNA levels are expressed as relative quantities in relation to the control strain. Both the control H. pylori 26695 mutant, possessing an intact arsRS locus, and an isogenic ΔarsS mutant were grown to mid-logarithmic stage of growth then harvested. **b-c)** After growth of the control H. pylori mutant and isogenic ΔarsS mutant to mid-logarithmic phase, equal aliquots were harvested and resuspended in pH 7 or pH 5 broth. mRNA levels are expressed as relative quantities in relation to the time zero sample. **b)** The expression of hsdM1 in the H. pylori 26695 control mutant at pH 7 or pH 5 at 30 and 60 minutes. **c)** The expression of hsdM1 in H. pylori ΔarsS at pH 7 or pH 5 at 30 and 60 minutes. Each dot represents a biological replicate, each done in a triplicate. Error bars; standard error of the mean. Statistical analysis via unpaired one-tailed t-test. *; P≤0.05, **; P≤0.01 ***; P≤0.001

Given that ArsRS is the major acid-sensing and response mechanism of *H. pylori,* and *hsdM*1 mRNA is impacted by the absence of ArsS, we hypothesized that *hsdM*1 expression is acid-responsive. We quantified *hsdM*1 mRNA in both the *H. pylori* 26695 control mutant and an *arsS* null mutant at pH 7 and pH 5. Our data indicate that *hsdM*1 transcription is induced at pH 5 in an ArsS-dependent manner (**Fig. 2b & c**).

The increased transcription of *hsdM*1 under pH 5 conditions peaks at 30 minutes. This may indicate that *hsdM*1 is part of an immediate acid response. However, the decrease in *hsdM*1 transcription at pH 5 at 60 minutes may be due to the induction of urease expression by pH 5 which raises the pH and thus may turn off the acid induction of *hsdM*1. *hsdM*1 is significantly downregulated at pH 5 in the *ΔarsS* mutant (**Fig. 2c**). We speculate that this may be due to the inability of this mutant to mount an acid acclimation response in the absence of a functional ArsRS TCS and thus its physiology is impaired by the continued low pH exposure.

*H. pylori* 26695 has two other complete Type I R-M systems with two Type I orphan specificity subunits **(Supplementary Table 1)**, and 16 Type II MTases, including orphan MTases and those within complete R-M systems. As part of our effort to examine the effects of environmental pH on the potential to methylate the *H. pylori* genome, and to determine if the acid response seen in *hsdM*1 was unique, we investigated four other MTases. We conducted qRT-PCR on the other Type I DNA MTases, *hsdM*2 (HP0850) and *hsdM*3 (HP1403), as well as two selected Type II DNA methyltransferases, *M.HpyAI* (HP1208) and *M.HpyAII* (HP1368). The results of our qRT-PCR analyses indicate that there are no significant changes in expression among any of these four DNA MTases in response to pH 5 conditions and that the ablation of the sensory histidine kinase ArsS has no apparent effect on these methyltransferases **(Supplementary Fig. 2-5)**. This indicates that the expression patterns revealed in the current study of *hsdM*1 are not universal among *H. pylori* restriction-modification systems or even among the Type I systems. The expression patterns seen in *M.HpyAI* and *M.HpyAII* **(Supplementary Fig. 4-5, respectively)** serve as a control to ensure that neither our *in vitro* acid shock itself nor the deletion of *arsS* are causing genome-wide changes in MTase expression. Therefore, ArsRS regulation and acid-sensitivity are unique to *hsdM*1 among Type I DNA methyltransferases, and possibly among other families of MTases, in *H. pylori* 26695.

### *hsdM*1 (HP0463) expression varies due to phase variation in the upstream gene, *hsdR*1 (HP0464)

A major means of bacterial acclimation to changing environments is the alternative expression states of operons and their resulting proteins. This process of phase variation is commonly the result of hyper-mutable repetitive DNA sequences that undergo slipped strand mispairing to alter the length of DNA repeats and thus turn on or off the expression of the affected gene^16^. A 2005 study by Salaun et al.^35^ identified six phase variable restriction-modification (R-M) systems in *H. pylori* 26695 and seven in the conspecific strain J99. Several of these phase variable loci were variably present among the strains examined in their study. In addition, they observed variation in the phase of these R-M systems among the studied strains. Other investigators have demonstrated that the phase variable nature of these R-M systems affect the expression of other genes, some related to virulence^36^. Gauntlett et al., 2014^31^ found that phase-locking *hsdR*1 (HP0464) in *H. pylori* OND79 had a detrimental effect on the colonization of mice.

We hypothesized that the length of the poly-C tract located within *hsdR*1 (HP0464), encoding the restriction endonuclease subunit of the *H. pylori* 26695 Type I R-M system HP0464-HP0463-HP0462, might affect the transcription of the downstream methyltransferase, *hsdM*1 (HP0463). Therefore, we created phase-stabilized versions of *hsdR*1 in *H. pylori* 26695. This was achieved by replacing the 3rd position cytosine of each proline codon in the poly-cytosine tract encoding prolines_205-209_ such that they still encoded tandem prolines, but slipped strand mispairing would be much less frequent during genome replication. This created a phase-lock ON allele of *hsdR*1. We created an alternate version of this stabilized *hsdR*1 that was phase-lock OFF by adding another cytosine after the stabilized proline codons such that HsdR1 could not be expressed as a full-length protein.

DNA sequencing of PCR amplicons from wild type *H. pylori* 26695 and both phase-locked ON and OFF *hsdR*1 mutants demonstrated that slipped strand mispairing, and thus the generation of variant *hsdR*1 alleles, was reduced greatly (data not shown). We subsequently used qRT-PCR to quantify *hsdM*1 mRNA in the *hsdR*1 phase-lock ON and phase-lock OFF *H. pylori* 26695 mutants and found an average 4-fold decrease in *hsdM*1 mRNA when the upstream gene in the *hsdR*1-*hsdM*1 operon was made phase-lock OFF (**Fig. 3**). Thus, within *H. pylori* populations, variants exist that express, and fail to express, this R-M system and may therefore possess differing degrees of genomic methylation.

**Figure 3.**
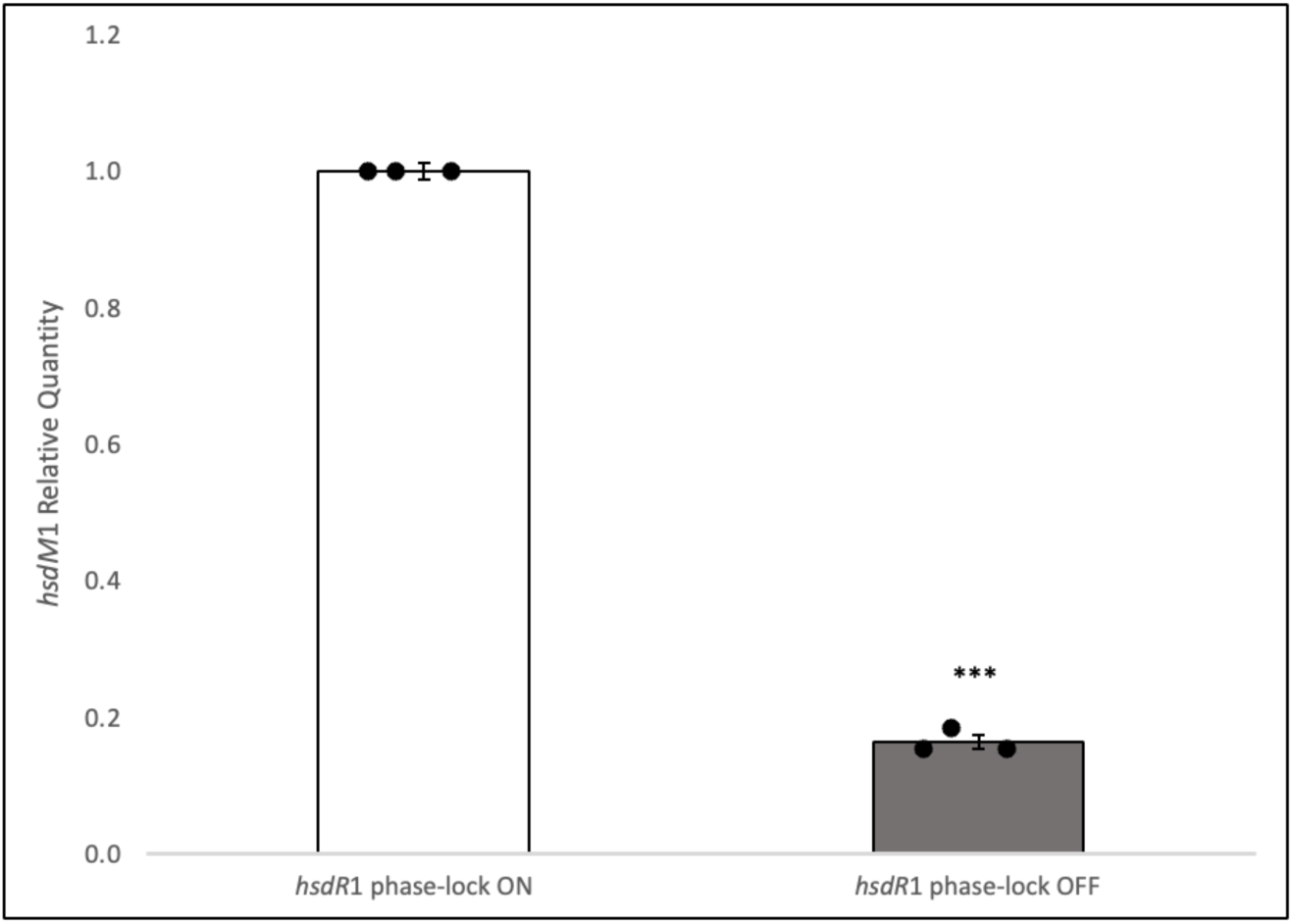
hsdM1 expression is significantly impacted by phase variation of hsdR1. hsdM1 mRNA levels in the hsdR1 phase-lock OFF mutant are expressed as relative quantities in relation to hsdM1 mRNA in the hsdR1 phase-lock ON mutant. Each dot represents a biological replicate, each done in triplicate. Error bars; standard error of the mean. Statistical analysis via unpaired one-tailed t-test. ***; P≤0.001

### Determination of the HsdM1 methylation motif

To characterize the impact of the HsdM1 methyltransferase on the epigenome, as well as the effect of acid and the ArsRS TCS, we cultured the *H. pylori* 26695 control mutant, isogenic *ΔarsS* mutant, and isogenic *ΔhsdM*1 mutant via serial passage in pH 7 or pH 5 liquid media, subculturing every 24 hours for three days. The *ΔhsdM*1 mutant allowed us to identify the target motif and regulatory impact of the Type I m6A DNA methyltransferase, HsdM1 (HP0463).

Type I restriction-modification systems recognize bipartite target sequences^22^. We determined the target motif of HsdM1 to be the complementary bipartite sequence pair 5’-(HH)TCA^m6A^VN_6_TGY-3’ and 3’-AGTN_6_GA^m6A^CA-5’. This determination is made because these are the only sequences that are methylated in the *H. pylori* 26695 control and not in the isogenic *ΔhsdM*1 mutants cultured under either pH condition **(Table 1**, **Table 2)**. This motif assignment has the same architecture as the target motif of HsdM2/M.HpyAXIII (HP0850), a previously characterized Type I m6A DNA methyltransferase in *H. pylori* 26695, which also has the ability to recognize and methylate its target motif on both DNA strands^37^. In the control *H. pylori* 26695 strain, 5’-(HH)TCA^m6A^VN_6_TGY-3’ is only methylated at pH 7 while 5’-(H)TCA^m6A^VN_6_TGY-3’ and 3’-AGTN_6_GA^m6A^CA-5’ are only methylated at pH 5 **(Table 2)**. 5’-(HH)TCA^m6A^VN_6_TGY-3’ is also not methylated in the *ΔarsS* strain under either pH 7 or pH 5 conditions **(Table 3)**. These data demonstrate that HsdM1 methylation is acid-dependent and that functionality is altered in the absence of *arsS.* This leaves some intriguing possibilities for the ability of HsdM1 to differentiate between unmethylated and hemi-methylated DNA, and the possible role of hemi-methylation in *H. pylori* epigenetic regulation.

**Table 1.**
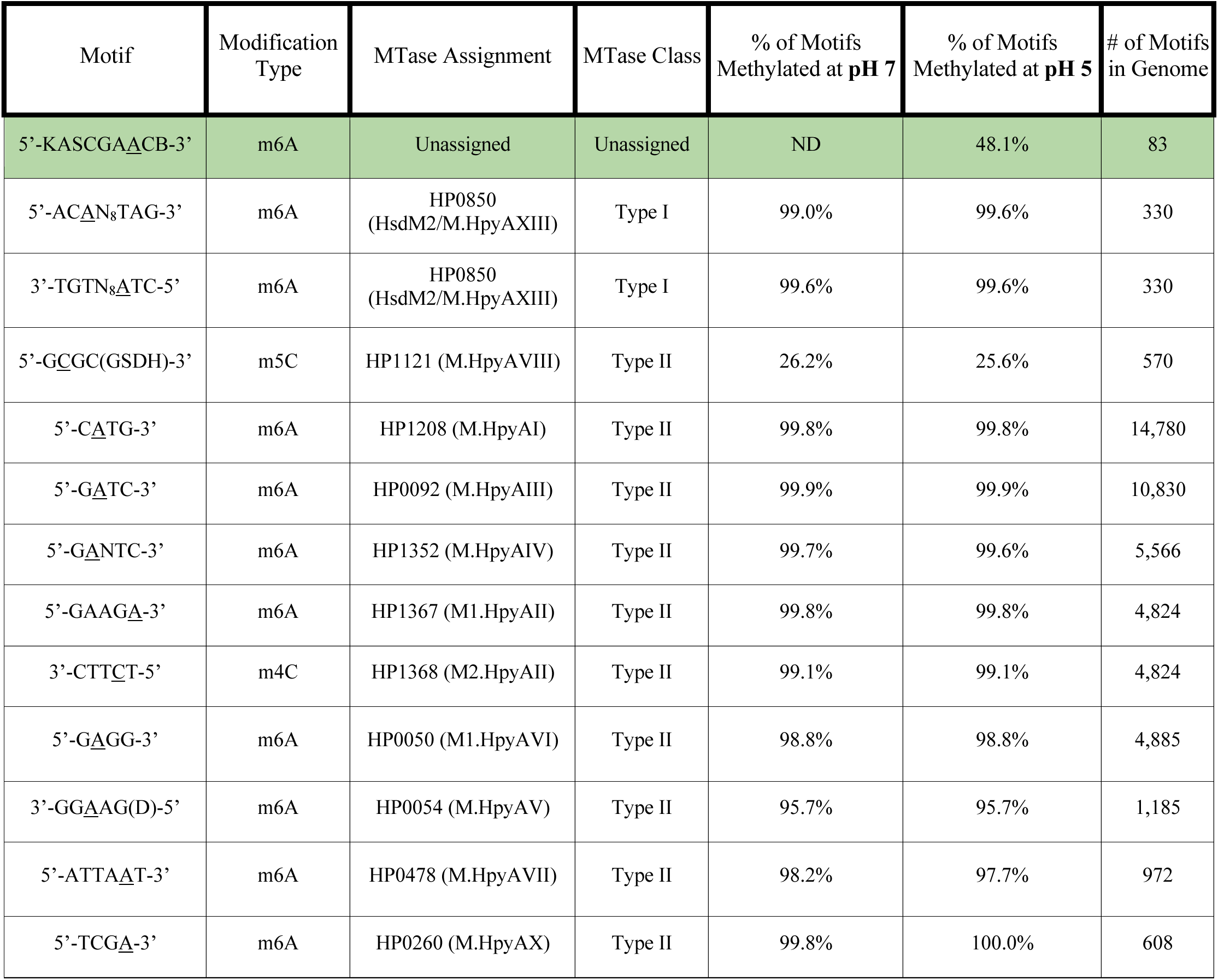
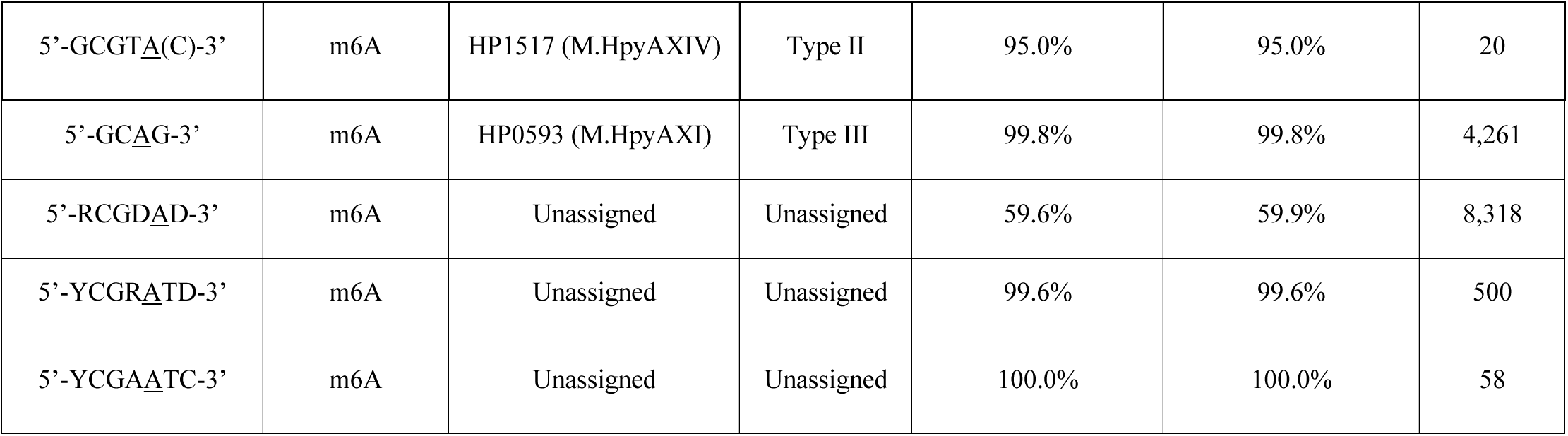
The methylomes of H. pylori 26695 ΔhsdM1 strain cultured at pH 7 or pH 5. Motif assignments are made based on Krebes et al., 2014^37^. Nucleotides in parenthesis are not part of the core motif. The samples were sequenced using PacBio Sequel SMRT Sequencing. The methylated nucleotide is underlined. Motifs highlighted in green are unique to a pH condition. ND; Not Detected.

**Table 2.**
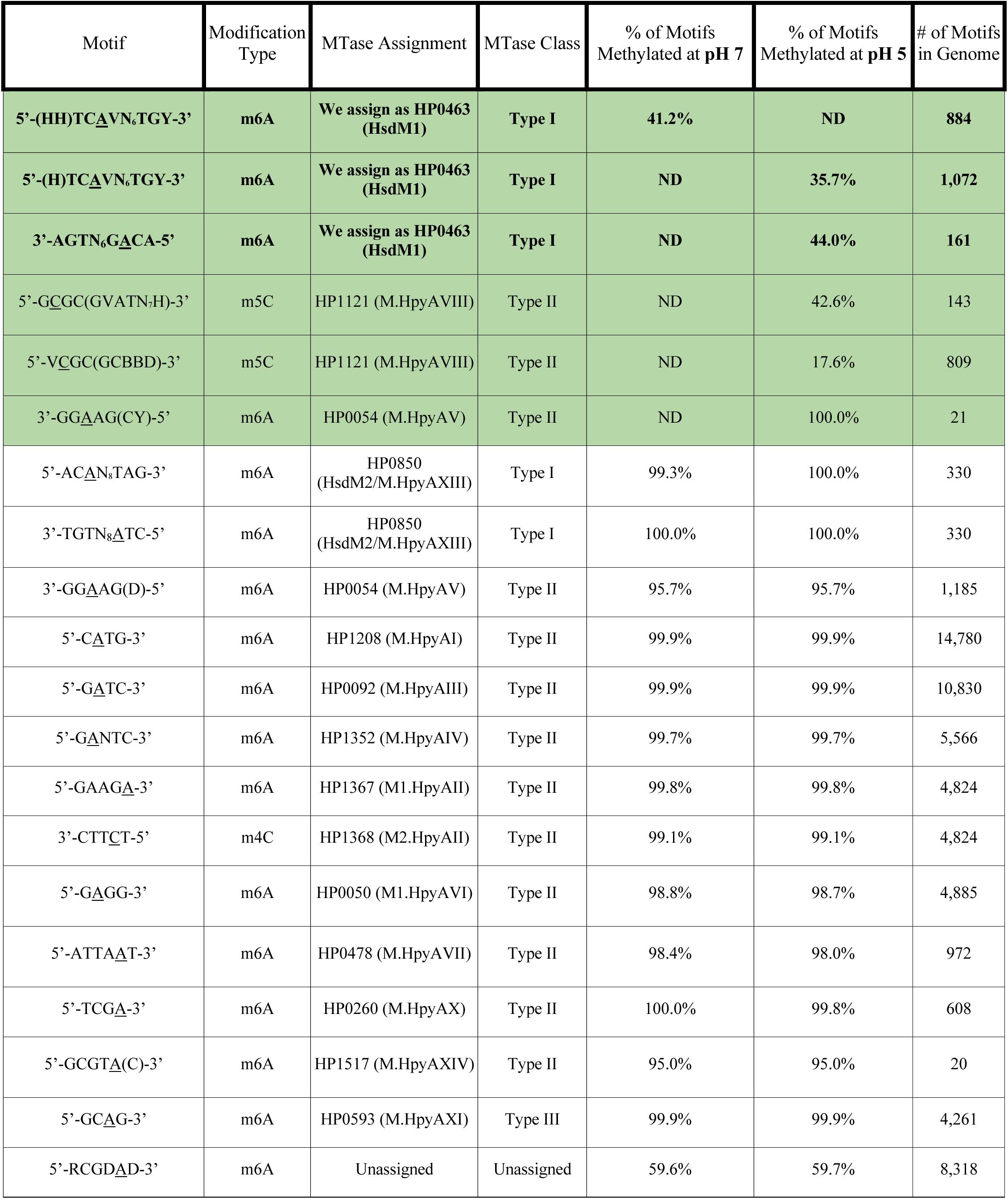
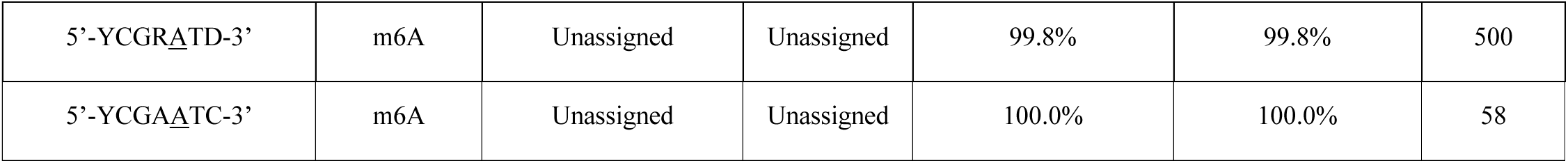
The methylomes of H. pylori 26695 control strain cultured at pH 7 or pH 5. Motif assignments are made based on Krebes et al., 2014^37^. Nucleotides in parenthesis are not part of the core motif. The samples were sequenced using PacBio Sequel SMRT Sequencing. The methylated nucleotide is underlined. Bolded motifs are assigned as the HsdM1 (HP0463) target motif. Motifs highlighted in green are unique to a single pH growth condition. ND; Not Detected.

**Table 3.**
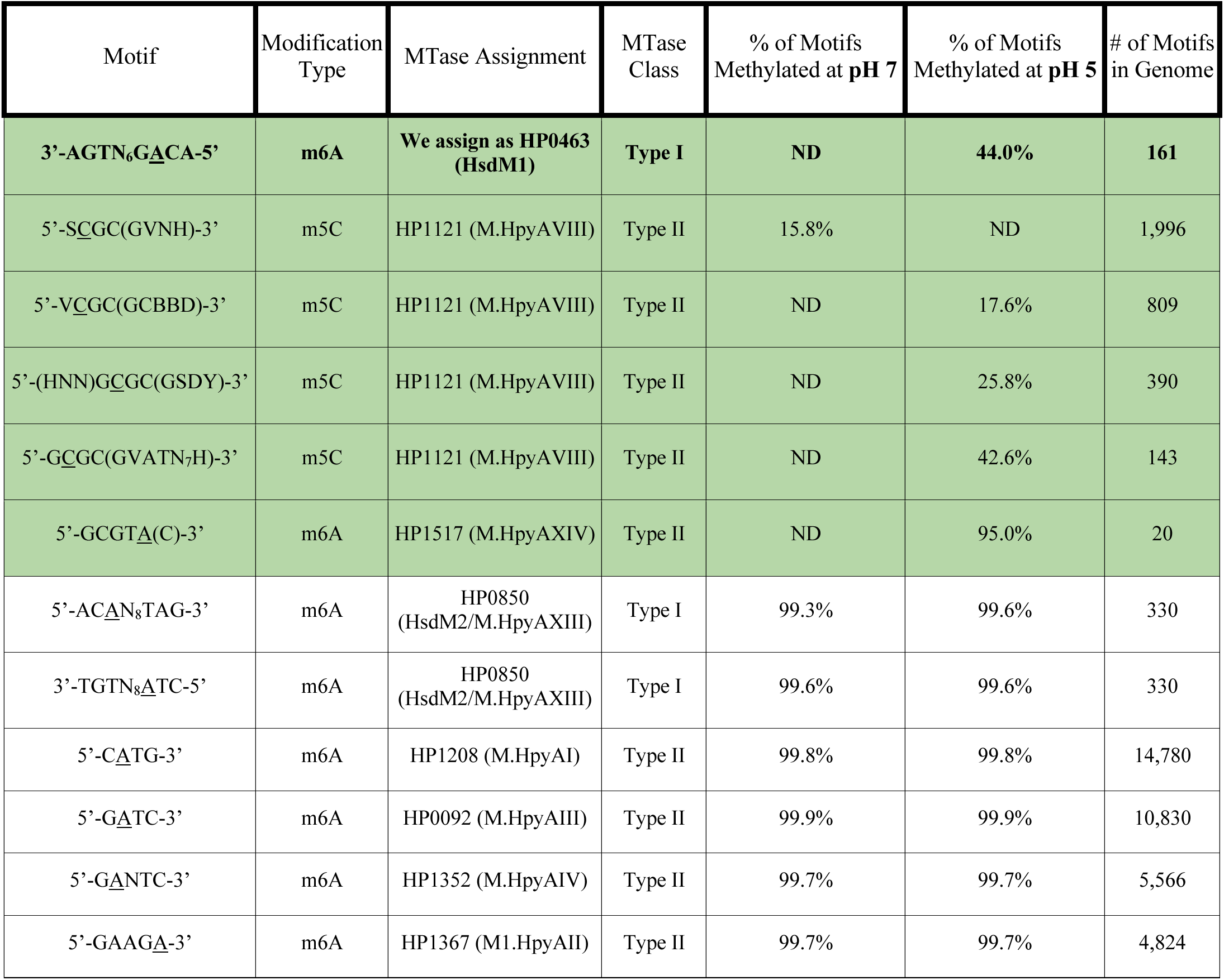
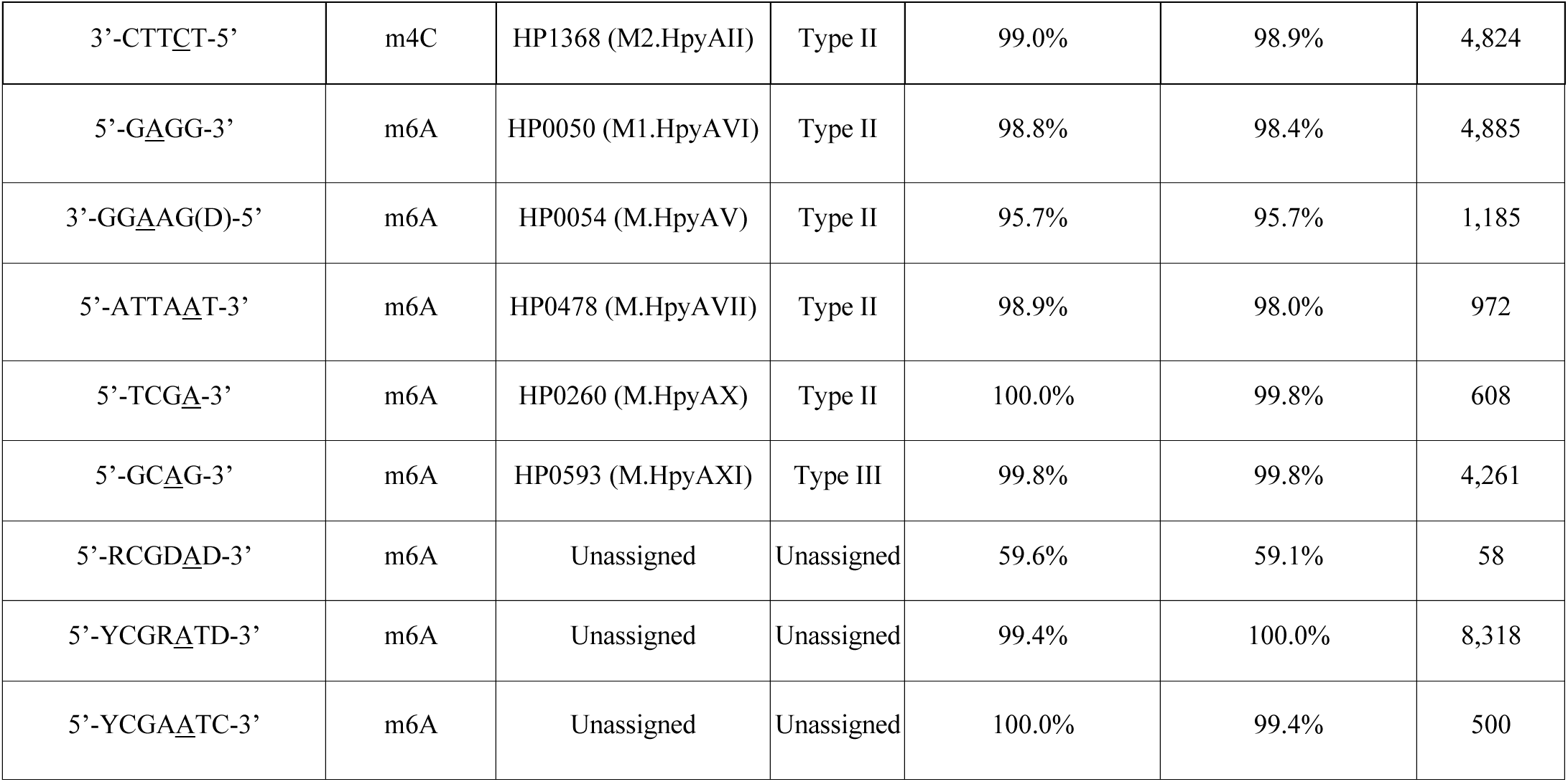
The methylomes of H. pylori 26695 ΔarsS strain cultured at pH 7 or pH 5. Motif assignments are made based on Krebes et al., 2014^37^. Nucleotides in parenthesis are not part of the core motif. The samples were sequenced using PacBio Sequel SMRT Sequencing. The methylated nucleotide is underlined. Bolded motifs are assigned as the HsdM1(HP0463) target motif. Motifs highlighted in green are unique to a single pH condition. ND; Not Detected.

The motif 5’-KASCGAA^m6A^CB-3 is methylated only at pH 5 in the *ΔhsdM*1 mutant. In addition, *ΔhsdM*1 is the only strain within which this motif is methylated. As 5’-CGAA^m6A^C-3’ are the nucleotides consistently called, we hypothesize that this is the core motif. This motif does not align with any currently annotated methyltransferases in *H. pylori* 26695, thus likely belongs to one of the many uncharacterized methyltransferases in *H. pylori* 26695.

There are multiple methylated motifs across our samples that PacBio Sequel Single Molecule, Real-Time (SMRT) Sequencing identified as including nucleotides that are not part of the established target motifs. In order to preserve the integrity of the methylome sequencing data without causing confusion, we have put these nucleotides in parentheses to indicate that they are not part of the accepted core motif. Interestingly, these extra nucleotides change between conditions for certain motifs, indicating that the genetic environment of the target motif may have an impact on methyltransferase activity. For example, the HsdM1 target motif on the forward strand is methylated at pH 7 when it possesses two leading H’s, but in *H. pylori* 26695 cells grown at pH 5, the methylated motif has only one leading H **(Table 2)**. M.HpyAV (HP0054) is a Type II m6A/m5C MTase with the m6A target sequence 3’-GGA^m6A^AG-5’^19,37^. In our samples, M.HpyAV (HP0054) methylates 3’-GGA^m6A^AG(D)-5’ at both pH 7 and pH 5 in *ΔhsdM*1, but methylates 3’-GGA^m6A^AG(CY)-5’ at pH 5 in the control **(Table 1**, **Table 2)**.

The variation in genetic environment also occurs with 5’-GC^m5C^GC-3’, the target motif of the Type II m5C MTase M.HpyAVIII (HP1121)^19,37,38^. Each sequenced strain in this study has 5’-GC^m5C^GC-3’ methylation occurring within different motif contexts. In the *H. pylori* 26695 control mutant, 5’-GC^m5C^GC(GVATN7H)-3’ and 5’-VC^m5C^GC(GCBBD)-3’ are methylated **(Table 2)**. In the *ΔhsdM*1 strain, only 5’-GC^m5C^GC(GSDH)-3’ is methylated **(Table 1)**. In the *ΔarsS* strain, 5’-SC^m5C^GC(GVNH)-3’, 5’-VC^m5C^GC(GCBBD)-3’, 5’-(HNN)GC^m5C^GC(GSDY)-3’, and 5’-GC^m5C^GC(GVATN7H)-3’ are methylated **(Table 3)**. Not only is there a loss of methylation of certain genetic environments in the mutants, but also additional methylated motifs upon the deletion of *arsS.* Therefore, we hypothesize that the varying nucleotides surrounding the core target motif may play a role in guiding selective methylation. Specifically, the variation in 5’-GC^m5C^GC-3’ genetic environments between the mutants indicates that M.HpyAVIII (HP1121) function is impacted by HsdM1 and ArsS.

Therefore, we come to the novel conclusions that the target motif of the Type I DNA methyltransferase HP0463 (HsdM1) is the complementary bipartite sequence pair 5’-(HH)TCA^m6A^VN_6_TGY-3’ and 3’-AGTN_6_GA^m6A^CA-5’, HsdM1 activity is acid-responsive in an ArsS-dependent manner, and selective methylation may be guided by the genetic environment of the target motif.

### The *H. pylori* methylome is acid-responsive

There are several studies characterizing both the transcription levels and protein dynamics of methyltransferases in acid conditions^9,39,40,41^. In addition, the *H. pylori* methylome in neutral conditions has been sequenced^37^. Here, we determine the *H. pylori* methylome in both neutral and acid growth conditions and demonstrate that the *H. pylori* methylome is acid-responsive.

In the *H. pylori* 26695 control, there are multiple sequences methylated at pH 5 and not at pH 7 **(Table 2)**. The 5’-GC^m5C^GC-3’ motifs, methylated by the Type II m5C DNA methyltransferase M.HpyAVIII (HP1121), and the bipartite sequence pair 5’-(H)TCA^m6A^VN_6_TGY-3’ and 3’-AGTN_6_GA^m6A^CA-5’, methylated by the Type I m6A DNA MTase HsdM1 (HP0463), are methylated in the genomes of the control mutant grown at pH 5, but not in pH 7 **(Table 2)**. As aforementioned, the M.HpyAVIII (HP1121) target motif exhibits two states of surrounding nucleotides, both of which are only methylated at pH 5, and the HsdM1 (HP0463) motif has an extra leading H when it is methylated at pH 7. Interestingly, while M.HpyAVIII (HP1121) motifs are not methylated in the *H. pylori* control grown at pH 7, these motifs are methylated to some degree in both the *ΔarsS* and *ΔhsdM*1 mutants grown at pH 7 **(Tables 1-3)**. Importantly, there are 2,206 additional instances of methylation in the genome of *H. pylori* 26695 when grown at pH 5 compared to the genome of the same strain grown at pH 7.

These data demonstrate for the first time that the *H. pylori* methylome is plastic and acid-responsive, and that the acid-responsiveness of the *H. pylori* methylome is partially dependent on HsdM1 (HP0463) and M.HpyAVIII (HP1121) activity.

### Methylation of the *H. pylori* genome is partially dependent upon the ArsRS TCS

When the ArsS acid-sensing histidine kinase of the ArsRS two-component system is deleted (*ΔarsS* strain), methylation of the HsdM1 forward target motif, 5’-TCA^m6A^VN_6_TGY-3’, is absent, while methylation of the complementary target motif, 3’-AGTN_6_GA^m6A^CA-5’, remains **(Table 3)**. This supports our earlier conclusion that *hsdM*1 expression is ArsS-dependent. This also indicates that HsdM1 may engage in hemi-methylation. The ability for methyltransferases to differentiate between unmethylated and hemimethylated DNA has previously been shown and could have exciting implications for the transcriptional and/or translational impacts^42^. Interestingly, our qRT-PCR data showed that *hsdM*1 expression is significantly upregulated at least 4-fold in *ΔarsS*, so it is surprising that HsdM1 activity appears to decline rather than increase. This suggests that post-transcriptional regulation may be occurring.

In the control strain, M.HpyAVIII (HP1121) motifs are only methylated at pH 5 **(Table 2)**. In *ΔarsS*, M.HpyAVIII (HP1121) motifs are methylated at both pH 5 and pH 7 with variations in the genetic environment **(Table 2)**. Both 5’-GC^m5C^GC-3’ motif contexts that are methylated in the *H. pylori* control mutant are also methylated in the isogenic *ΔarsS* mutant, but two additional genetic environments are methylated in the genome of the *ΔarsS* mutant. Of particular interest, 5’-SC^m5C^GC(GVNH)-3’ is unique to the *ΔarsS* mutant and is present 1,996 times in the genome which is over 1,000 more occurrences than any other 5’-GC^m5C^GC-3’ motif environment in any strain **(Table 3)**. It’s also the only 5’-GC^m5C^GC-3’ motif that is methylated in pH 7 conditions in the *ΔarsS* mutant **(Table 3)**. This supports our hypothesis that the genomic environment of target motifs plays a role in selective methylation. Therefore, we conclude that M.HpyAVIII (HP1121) is under the regulation of ArsRS and this regulation impacts the methylome.

M.HpyAXIV (HP1517) is active in the *H. pylori* 26695 control and *ΔhsdM*1 isogenic mutant grown at both pH 7 and pH 5 conditions **(Table 1 and Table 2)**, suggesting that the expression of this N6 adenosine methyltransferase is independent of pH or the hierarchical influence of HsdM1. However, the M.HpyAXIV target motif is methylated in the *ΔarsS* mutant only when grown at pH 5 **(Table 3)**. This suggests the expression of M.HpyAXIV in *H. pylori* grown at pH 7 may be affected by the ArsRS TCS.

## Discussion

It is becoming increasingly clear that restriction-modification (R-M) systems, particularly DNA methyltransferases, play an important role in gene regulation beyond their role in genome protection. *Helicobacter pylori* is an important organism in which to study the different functions of methyltransferases as it possesses over 20 R-M systems, a significant over-representation for its small genome of 1.64–1.67 Mb^19, 26^. Our current study demonstrates regulatory mechanisms and patterns of the Type I DNA methyltransferase HsdM1 (HP0463) as well as the role of phase variation, acid, and the acid acclimation facilitator, the two-component system (TCS) ArsRS in *H. pylori* epigenetics. Our data demonstrate that *hsdM*1 expression is induced under acidic conditions *in vitro* and that this acid response is ArsS-dependent.

The methylated DNA motifs of interest determined in the current study have comparatively low abundances and varying degrees of methylation compared to most of the annotated motifs. The most ubiquitous motifs in the genome, such as 5’-CA^m6A^TG-3’ which appears 14,780 times and 5’-GA^m6A^TC-3’ which appears 10,830 times, are both methylated in all our samples to at least 98.8% methylation. This may indicate that these motifs play a more traditional genome protection role. The intermediate levels of methylation of the HsdM1 (HP0463) target motifs, between 35.7% and 44.0% methylation, and the M.HpyAVIII (HP1121) target motifs, between 17.6% and 42.6% methylation, challenge the idea that methylation is an on/off switch and supports the growing hypothesis that certain methyltransferases have a primarily regulatory role^30^.

Existing studies of the *H. pylori* methylome focus only on the core motifs of the methyltransferases when characterizing methyltransferase activity. However, we speculate that the story may be much more complex. In this study, we included all nucleotides that PacBio SMRT Sequencing annotated as part of each methylated motif, even if they are not part of the annotated core motifs of previous studies. We found that the genetic environment of certain methylated motifs changed between pH 7 and pH 5 conditions and among our three *H. pylori* mutants. Therefore, we make the novel proposal that the genetic environment of the methyltransferase core DNA motifs may play a role in guiding selective methylation, and thus could mediate differential selective methylation in varying conditions.

The expression patterns of *hsdM*1 do not appear to follow the traditional model of two-component system (TCS) regulation. In neutral pH conditions, the standard view of ArsRS TCS functioning predicts a high quantity of unphosphorylated response regulator, ArsR, and a low quantity of phosphorylated ArsR (ArsR∼P)^10^. The sensory histidine kinase protein ArsS adopts different conformations based on the level of acidity via varying degrees of protonation of the signal input domain, thus producing varying concentrations of ArsR∼P^10^. In the traditional model, response regulators are dimerized and act as transcription factors when phosphorylated^9^. This model predicts that as ArsR represses *hsdM*1 expression under neutral conditions *in vitro,* as evidenced by the de-repression of *hsdM*1 expression in our *ΔarsS* mutant (**Fig. 2a**). Therefore, we would expect *hsdM*1 to be repressed in acidic conditions as there is a substantially higher concentration of ArsR∼P predicted due to activated ArsS. However, this simple model is not sufficient to explain our demonstration that acidic pH induces *hsdM*1 transcription in an ArsS-dependent manner, rather than repressing it. This leads us to speculate that non-phosphorylated ArsR may also be acting as a transcriptional regulator of *hsdM*1. Non-phosphorylated ArsR has been found to act as a transcriptional regulator, and *ΔarsR* strains are not viable, which indicates that ArsR in the non-phosphorylated form has an essential role within *H. pylori*^9,43^. It is tempting to speculate that ArsR and ArsR∼P may bind the same DNA sites with different affinities, or that the manner in which ArsR and ArsR∼P bind the DNA results in different DNA conformations and thus different transcriptional outcomes.

Preliminary analysis of the genomic locations of methylations reveals the presence of differentially modified bases in certain promoter regions. The antisense promoter region of M.HpyAVIII (HP1121), annotated by Sharma et al., 2010^34^, contains a modified adenine nucleotide in *H. pylori* strain *ΔhsdM1* grown at pH 7 and pH 5 conditions and *ΔarsS* at pH 5 conditions. This supports our hypothesis that M.HpyAVIII (HP1121) expression is regulated by HsdM1 and ArsS. Additionally, HP0731, a gene involved in toxin production, as annotated by Kato et al., 2017^44^, contains a methylated 5’-(H)TCA^m6A^VN_6_TGY-3’ motif, the target of the methyltransferase HsdM1 (HP0463), within one of the antisense promoters identified by Sharma et al., 2010^34^, in the *H. pylori* 26695 control strain grown at both pH 5 and pH 7 conditions. The presence of methylation in promoter regions supports the hypothesis that methylation plays a regulatory role.

Lastly, we noticed a modified guanine nucleotide within the primary promoter of the HP0052-HP0056 operon in the *H. pylori* 26695 control and *ΔarsS* strains at both pH conditions. This guanine methylation is likely an oxidative modification, rather than due to the action of a methyltransferase^45^. While 8-oxoG modifications can, in some cases, cause lesions preventing binding of DNA and RNA polymerase, we speculate a potential role HsdM1 may play in regulating transcription of an 8-oxoG repair system, as the guanine base modification was present in high enough frequency to be reported in both the control and *ΔarsS* samples, but not in the *ΔhsdM*1 samples^46^. These results suggest that any change in M.HpyAV (HP0054) transcription by *hsdM1* is indirect, and *hsdM1* may instead be acting upon a hypothetical 8-oxoG repair system.

This possible regulation is fascinating as HP0054 encodes the Type II m6A/m5C MTase M.HpyAV, with the target motifs 5’-CC^m5C^TTC-3’ and 3’-GGA^m6A^AG-5’^19,37^. M.HpyAV (HP0054) is the only methyltransferase in *H. pylori* 26695 that can be both an m6A and m5C MTase^19,37^. However, it exhibits only m6A activity in our samples. In addition, the gene directly upstream of this guanine modification is M2.HpyAVI (HP0051), a Type II m5C MTase, with the target motif 5’-C^m5C^CTC-3’^19,37,38^. Neither of these m5C motifs are methylated in any of our samples, leaving M.HpyAVIII (HP1121) as the only active m5C methyltransferase in our samples. The possible regulation of the hypothetical 8-oxoG repair system and m5C methyltransferases adds another layer to the web of complex, hierarchical methyltransferase regulation.

Interestingly, there are some discrepancies between our data and those of some existing studies. For example, Banajaree & Rao, 2011^40^ and Narayanan et al., 2020^41^ both found that M.HpyAXI (HP0593) has an acid optimum at pH 5.5 in solution and can only form a functional tetramer in acidic conditions. RNA sequencing data from Wen et al., 2003^39^ also found M.HpyAXI (HP0593) to be transcriptionally upregulated 2.5-fold in acid conditions. Therefore, we would expect its target motif, 5’-GCA^m6A^G-3’, to have significantly higher methylation at pH 5. However, our methylome sequencing revealed an equal percentage of methylation of the M.HpyAXI (HP0593) target motif, 5’-GCA^m6A^G-3’, in pH 7 and pH 5 in all strains **(Tables 1-3)**, thus indicating M.HpyAXI (HP0593) enzymatic activity *in vivo* is unaffected by acidic pH. This may be because, as the gastric pH drastically fluctuates, *H. pylori* has mechanisms for intracellular buffering, such as the urease gene cluster^39,47^. Therefore, despite an extracellular pH of 5, the intracellular pH likely does not drop to pH 5. Thus, the Banajaree & Rao, 2011^40^ and Narayanan et al., 2020^41^ studies of protein dynamics *in vitro* may not translate *in vivo*, at least not in non-extreme extracellular pHs.

Wen et al., 2003^38^ also demonstrates transcriptional upregulation of M.HpyAIV (HP1352), M.HpyAVII (HP0478), and M.HpyAI (HP1208) in acidic conditions. None of the target motifs of these methyltransferases exhibit a differential percentage of methylated motifs at pH 7 vs pH 5 in any of our samples.

Discrepancies between predicted methylation based on transcriptional data and actual methylation patterns, both within this study and in comparison to previous studies, supports the growing hypothesis of complex, layered epigenetic regulation. There are likely complex regulation mechanisms that manifest at the translational and/or protein levels. For example, hierarchical regulation between *H. pylori* DNA methyltransferases via methylation at translational start and stop codons has been previously demonstrated^27^. This may help explain the significant variance between studies characterizing the ArsRS regulon^2,9,11,12,39^. Of particular interest, as we identified differential methylation in antisense promoters, antisense RNA has been shown to regulate R-M systems in *E. coli*^48^. For Dam in *E. coli*, switching between fully methylated and hemimethylated DNA has been shown to directly regulate phase variation systems^49,50^.

Yano et al., 2020^27^ demonstrated that HsdM1 is hierarchically regulated in *H. pylori* 22695 by M.HpyAXI (HP0593), HsdM2/M.HpyAXIII (HP0850), and M.HpyAX (HP0260). However, they did not investigate whether HsdM1 regulates any other methyltransferases. We are making the novel suggestion that the Type II m5C DNA methyltransferase M.HpyAVIII (HP1121) plays a key role in acid response in the methylome and is regulated by the two-component system ArsRS and likely via hierarchical regulation by HsdM1 (HP0463). In the *H. pylori* 26695 control strain, M.HpyAVIII (HP1121) is active at pH 5 conditions, but not at pH 7 conditions **(Table 2)**. However, M.HpyAVIII (HP1121) is active in both pH conditions in the *ΔhsdM1* and *ΔarsS* samples **(Table 1)**. In addition, the antisense promoter of M.HpyAVIII (HP1121) is methylated in *ΔarsS* at pH 5 and *ΔhsdM*1 at pH 5 and pH 7, but never in the control. Therefore, this m6A methylation is occurring as a byproduct of either the loss of regulation or absence of HsdM1. This demonstrates that the functionality of M.HpyAVIII (HP1121) is sensitive to the presence of HsdM1 and ArsS. This is interesting as M.HpyAVIII (HP1121) appears to be the only active m5C DNA methyltransferase in *H. pylori* 26695, and thus may play a unique role in gene regulation.

It is worth noting that the PacBio SMRT Sequel technology annotated 5’-GC^m5C^GC-3’ as m4C methylation, but we attribute the 5’-GC^m5C^GC-3’ methylation to the Type II m5C DNA methyltransferase M.HpyAVIII (HP1121) as this specificity has been previously demonstrated^19,37,38^. *H. pylori* 26695 has been shown multiple times to only have one m4C MTase, M2.HpyAII (HP1368), with the target motif 3’-CTTC^m4C^T-5’^20,37,38^ which was correctly identified in our data. The PacBio Sequel machine does not have the capacity to differentiate between m4C and m5C methylation, as they are too chemically similar, and has been proven to mistakenly annotate m5C methylation as another type, particularly in *H. pylori* ^51,52^.

We show, for the first time, that a two-component system can indirectly impact gene expression by demonstrating that differential transcriptional activation of methyltransferase genes has an impact on the epigenome, and thus likely on the transcription of other genes. We characterize the target motif of the Type I m6A DNA methyltransferase HsdM1 (HP0463) and demonstrate that both its transcription and protein activity, as shown by differential methylation patterns of its target motif, are regulated by the two-component system ArsRS. We show that the Type II m5C DNA methyltransferase M.HpyAVIII (HP1121) is regulated by ArsRS, and likely under the hierarchical control of HsdM1. In addition, we propose that the genetic environment of methyltransferase target motifs play a role in guiding selective methylation. These methyltransferase dynamics result in an acid-sensitive, plastic epigenome in *H. pylori* 26695. This study expands the possibilities for *H. pylori*’s ability to sense and respond to the dynamic environment of the human gastric epithelium which may play a key role in the decades-long persistent infection by *H. pylori*.

## Material and Methods

### Culture conditions

*H. pylori* 26695 strains were grown on Trypticase Soy Agar II plates with 5% sheep blood and in Sulfite-Free Brucella Broth containing cholesterol (Gibco-BRL) (SFBB-chol) at 37°C in 5% CO_2_*/* 95% ambient air.

### Bacterial mutants

*H. pylori* strain 26695 *ΔrdxA*, a metronidazole resistant mutant, served as the control in all experiments and is referred to hereafter as the control mutant. The *ΔrdxA* mutant was used as part of a counter-selection procedure to create the *ΔhsdM*1 mutant described below as well as the *ΔarsS* mutant. The counter-selection procedure is detailed in Loh et al., 2011^53^ and the *ΔarsS* construction procedure is detailed in Loh et al. 2021^12^. This controls for any potential impact of the *ΔrdxA* locus. This locus is unrelated to any studied functions as it encodes an oxygen-insensitive NADPH nitro-reductase and its deletion confers metronidazole resistance.

*H. pylori* 26695 *ΔrdxA-ΔhsdM*1, referred to hereafter as *ΔhsdM*1, is the deletion of *hsdM*1 (locus tag; HP0463). A 2081 bp fragment of *H. pylori* strain 26695, containing the coding sequence of *hsdM*1 (HP0463), was amplified by PCR using; 5’-CACGAACAAACGCCTTCATAAGCACC-3’ and 5’-TAAGCGTGAAATGCTGCGGCTGCC-3’. The amplicon cloned into pGEM T-EASY (Promega). An 804 bp portion of the *hsdM*1 coding sequence was deleted by inverse PCR with the inclusion of a unique *Bam*HI site in each of the following primers; 5’-GGG***<u>GGATCC</u>***GGCGAGTTTGGGGAGTCTTTTATCGG-3’ and 5’-GGG***<u>GGATCC</u>***TGCTTAGTAATAAGGGTAAGGGGGC-3’. Introduced *Bam*HI sites are indicated in bold italics and underlined. After amplification and *Bam*HI digestion, the CAT/*rdxA* fragment encoding from pMM674^53^ was cloned into the introduced *Bam*HI site. Natural transformation of this plasmid, containing a deletion within *hsdM*1 along with an intact copy of *H. pylori rdxA* (HP0954) and the gene for chloramphenicol resistance, into *H. pylori* 26695 *ΔrdxA* was used for allelic replacement of the native *hsdM*1 gene. Subsequent counter-selection by the method of Loh et al. 2011^53^ was used to introduce a markerless 804 bp deletion allele of *hsdM*1. Sequence was confirmed by PCR amplification and sequencing.

*H. pylori* 26695 *hsdR*1 phase-lock variants were created. HsdR1 (HP0464) is a predicted Type I restriction endonuclease. The region of the gene encoding amino acids 205-210 (TPPPPQ) possesses a 15 nt poly-cytosine tract that we hypothesized facilitates phase variation of *hsdR*1 via Slipped Strand Mispairing (SSM) resulting in an increased mutation rate creating sub-populations of *H. pylori* 26695 with *hsdR*1 phase-lock OFF and phase-lock ON. To limit or prevent this region from undergoing SSM, *hsdM*1 sequences were synthesized (GeneWiz) in which the 3rd position of each of the 4 proline codons (206-209) was changed to a T, A, or G (CCT_206_ CCA_207_ CCG_208_ CCT_209_). This preserved the proline coding capacity while greatly reducing the mutation rate. Another version of this sequence synthesized possessed an additional C placed after the modified proline codon 209 to create a phase-lock OFF version of *hsdR*1. Both versions of *hsdR*1, phase-locked ON and phase-locked OFF, were cloned into pUC-GW-Amp. To facilitate screening of allelic replacement of these mutant forms of *hsdR*1 into *H. pylori* 26695, a mutant version of each of the *hsdR*1 had a unique *Bam*HI site inserted by site directed mutagenesis of a C at position 1033 of *hsdR*1 to a G using oligonucleotide directed mutagenesis (5’-CATCAGAGACTTTTTTAGC***<u>G</u>***<u>GATCC</u>AACCTAAACAAAAAGAC-3’) (Quick-Change Lightning. Agilent). The mutated site is shown in bold and italics and the generated *Bam*HI site is underlined. A CAT-*rdxA* gene cassette was cloned into this *Bam*HI site from pMM674^53^, a gift of Drs. Mark McClain and Timothy Cover, Vanderbilt University. The chloramphenicol and metronidazole counter-selection of Loh et al., 2011^53^ was used to place each of the *hsdR*1 phase-locked ON and phase-locked OFF gene cassettes into *H. pylori* 26695 and was confirmed by PCR and sequencing.

### RNA extraction

Upon collection, samples were centrifuged at 6,000 x g for 7 minutes then resuspended in RNAzol RT (Molecular Research Center). Samples then underwent RNA extraction. The RNA was quantified for concentration and purity using a Nanodrop (ThermoFisher), and the purified RNA samples were stored at −80°C.

### Quantitative Real-Time Polymerase Chain Reaction (qRT-PCR)

All samples that underwent RNA extraction were used as templates for the synthesis of cDNA for Quantitative Real-Time PCR (qRT-PCR). cDNA synthesis (iScript, Bio-Rad) was performed as specified by the manufacturer (Bio-Rad). TaqMan custom gene expression assays for the Type I MTases *hsdM*1 (HP0463), *hsdM*2 (HP0850), and *hsdM*3 (HP1403), and the Type II MTases *M.HpyAI* (HP1208) and *M.HpyAII* (HP1368) were synthesized by ThermoFisher. The sequences of all probes and reporters used can be found in **Supplementary Table 2**. All reporters have FAM as the fluorophore at the 5’ end and NQR as the quencher at the 3’ end. qRT-PCR was performed on the StepOne Plus System or QuantStudio3 System (Thermo Fisher). Target and normalizing genes in each sample were analyzed in technical triplicates. qRT-PCR data was analyzed by calculating Relative Quantity (RQ) values against GyrB using the 2^-ΔΔCT^ method^54^. Due to the extreme sensitivity of qPCR, technical replicates with an RQ standard deviation over 0.25 were considered technical errors and were not included^54^. The DNA Gyrase B subunit gene (*gyrB*) was used as the normalizing gene as previously described^55,56^.

### Acid shock experiments

*H. pylori* 26695 control mutant and the isogenic Δ*arsS* mutant were subjected to acid shock experiments at pH 5 for 60 minutes. *H. pylori* cells were harvested from plates and resuspended in 2.5 ml of Sulfite-Free Brucella Broth containing cholesterol (Gibco-BRL) (SFBB-chol) at pH 7. These mutants were cultured in an incubator at 37° C in a 5% CO_2_*/* 95% ambient air environment with shaking at 150 rpm. The growth of the two overnight cultures was quantified via OD_600_. New broth cultures were begun by subculturing each overnight culture to achieve a final OD_600_ of 0.4. These cultures were incubated for 7 hours shaking in a 5% CO_2_ incubator at 37° C under the conditions described before.

After 7 hours, the OD_600_ of the two cultures were taken and 1 OD_600_ unit (∼10^9^ cells) from the control and Δ*arsS* mutant cultures were collected as the time zero samples. The control and Δ*arsS* mutant cultures were each split into two equal aliquots and harvested by centrifugation at 5,000 x g, 20°C, for 10 min. The supernatant media was decanted, and one of the two pellets of each mutant *H. pylori* was resuspended in the same volume of SFBB-chol at pH 7, while the other pellet was resuspended in the same volume of media, but at pH 5, and then both were transferred to a six-well plate. The plate was incubated for 60 minutes under the same conditions described for broth cultures above with samples taken at 30 minutes and 60 minutes. To collect the *H. pylori* cells, 1 OD_600_ unit of the cultures was harvested by centrifugation at 6,000 x g. The supernatant was decanted, cells were resuspended in 1mL RNAzol (Sigma-Aldrich), and samples were stored at −80°C. All samples were subsequently subjected to RNA extraction and cDNA synthesis in preparation for qRT-PCR to determine the concentrations of mRNA of interest.

### Acid growth experiment

We conducted a three-day, serial passage acid growth experiment using the *H. pylori* 26695 control mutant and isogenic *ΔarsS* and *ΔhsdM*1 mutants. All strains were initially grown on Trypticase Soy Agar/5% sheep’s blood plates. Overnight broth cultures and subcultures were performed as in the acid shock experiments. After 7 hours of subculture, each strain was split into two equal aliquots and *H. pylori* cells were isolated by centrifugation at 5,000 x g, 20° C, for 10 min. The supernatant media was decanted, and one of the two pellets of each *H. pylori* mutant was resuspended in SFBB-cholesterol-vancomycin at pH 7, while the other pellet was resuspended in the same media at pH 5 and transferred to a six-well plate. These cultures were grown overnight under standard broth culture conditions. The next morning, each culture was subcultured by transferring 0.2 OD_600_ units of each culture into 6 mL of new media to make a new overnight culture. This subculture procedure was repeated the next morning, and samples were collected the morning after that so that each mutant was growing in pH 5 or pH 7 broth for 72 hours total. Culture was harvested by centrifugation at 4,500 rpm, 20°C, for 10 min. The supernatant was decanted, and the pellets were stored at −20°C. The samples were then prepped for methylome sequencing, described below.

### Methylome sequencing

Samples were prepared via genomic DNA extraction using the Bio-Rad genomic DNA extraction kit via the manufacturer’s suggested protocol including the optional RNA elimination step. The extracted DNA was quantified for concentration and purity using a Nanodrop (Thermo Fisher). As an additional quality check, we performed agarose gel electrophoresis to ensure all extracted DNA was above 5kB and had not been fragmented.

The samples were sequenced by Azenta Life Sciences using the PacBio Sequel technology. This determines N^6^-methyladenosine and N^4^-methylcytosine methylation status genome-wide and identifies methylation motifs using Single Molecule, Real-Time (SMRT) sequencing.

### Statistical analysis

Significant differences between the means of the control and treatment groups were analyzed via one-tailed unpaired t-test. Data were expressed as the means ± standard errors of the means.

## Data Availability

The SMRT Sequence data discussed in this publication have been deposited in NCBI’s Gene Expression Omnibus and are accessible through GEO Series accession number GSE241991.

## Acknowledgements

Research was funded through a grant from The National Institutes of Health, NIH-R15 A1133470 (MHF).

