## supplementary figures for "The *Helicobacter pylori* Methylome is Acid-Responsive due to Regulation by the Two-Component System ArsRS and the Type I DNA Methyltransferase HsdM1 (HP0463)"

| **Locus Tag** | **Gene Name** | **Protein Type** |
| --- | --- | --- |
| HP0464 | *hsdR*1 | Restriction Endonuclease |
| HP0463 | *hsdM*1 | DNA Methyltransferase |
| HP0462 | *hsdS*1b | Specificity Subunit |
| HP0846 | *hsdR*2 | Restriction Endonuclease |
| HP0850 | *hsdM*2 | DNA Methyltransferase |
| HP0848 | *hsdS*2 | Specificity Subunit |
| HP1402 | *hsdR*3 | Restriction Endonuclease |
| HP1403 | *hsdM*3 | DNA Methyltransferase |
| HP1404 | *hsdS*3b | Specificity Subunit |
| HP0790 | *hsdS*5 | Specificity Subunit |
| HP1383 | *hsdS*6 | Specificity Subunit |

**Supplementary Table 1.** *H. pylori 26695 Type I Restriction-Modification Systems.*

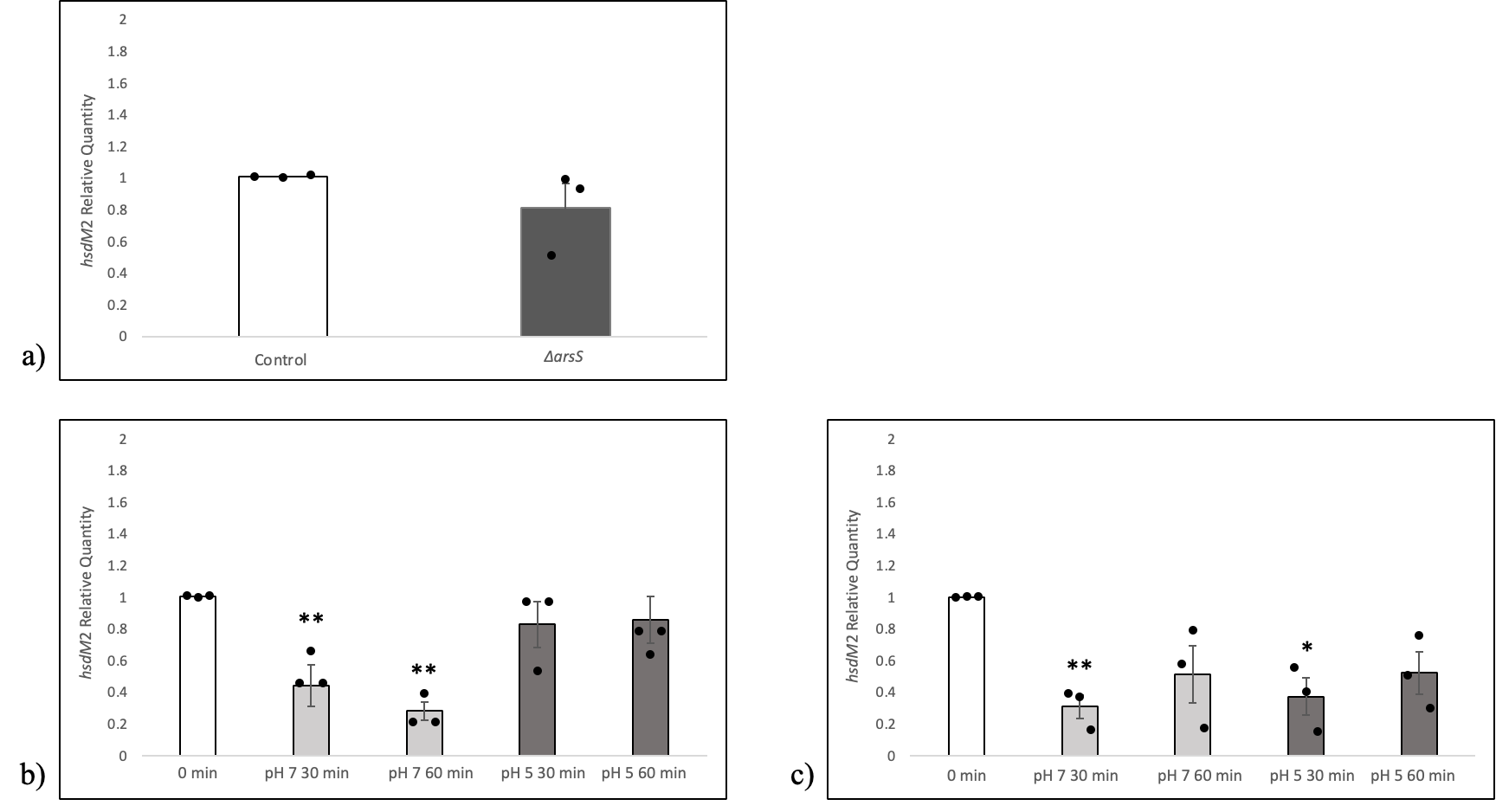

**Supplementary Figure 2. *hsdM*2 (HP0850) transcription is neither acid-sensitive nor ArsS-dependent.** *Both the control H. pylori mutant, possessing an intact arsRS locus, and an isogenic ΔarsS mutant were grown to mid-logarithmic stage of growth and equal aliquots were harvested and resuspended in pH 7 or pH 5 broth.* **a)** *hsdM2 mRNA levels are expressed as relative quantities in relation to the control strain. Both the control H. pylori mutant, possessing an intact arsRS locus, and an isogenic ΔarsS mutant were grown to mid-logarithmic stage of growth and equal aliquots then harvested.* ***b-c)*** *After growth of the control H. pylori mutant and isogenic ΔarsS mutant to mid-logarithmic phase, equal aliquots were harvested and resuspended in pH 7 or pH 5 broth. mRNA levels are expressed as relative quantities in relation to the time zero sample.* ***b)*** *The expression of hsdM2 in the H. pylori 26695 control mutant at pH 7 or pH 5 at 30 and 60 minutes.* ***c)*** *The expression of hsdM2 in H. pylori ΔarsS at pH 7 or pH 5 at 30 and 60 minutes. Each dot represents a biological replicate, each done in a triplicate. Error bars; standard error of the mean. Statistical analysis via unpaired one-tailed t-test. *; P≤0.05, **; P≤0.01*

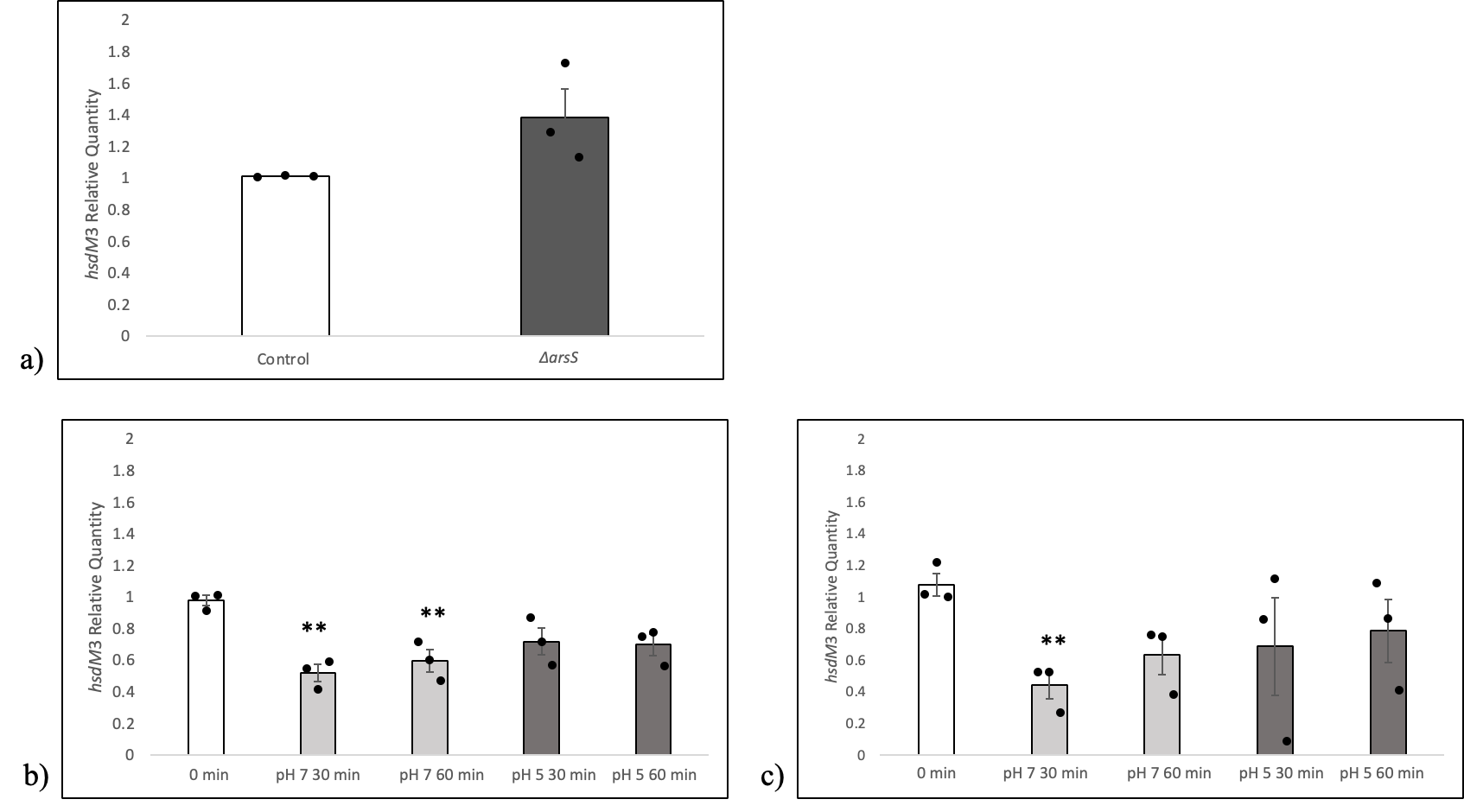

**Supplementary Figure 3. *hsdM3* (HP1403) transcription is neither acid-sensitive nor ArsS-dependent.** *Both the control H. pylori mutant, possessing an intact arsRS locus, and an isogenic ΔarsS mutant were grown to mid-logarithmic stage of growth and equal aliquots were harvested and resuspended in pH 7 or pH 5 broth.* **a)** *hsdM3 mRNA levels are expressed as relative quantities in relation to the control strain. Both the control H. pylori mutant, possessing an intact arsRS locus, and an isogenic ΔarsS mutant were grown to mid-logarithmic stage of growth and equal aliquots then harvested.* ***b-c)*** *After growth of the control H. pylori mutant and isogenic ΔarsS mutant to mid-logarithmic phase, equal aliquots were harvested and resuspended in pH 7 or pH 5 broth. mRNA levels are expressed as relative quantities in relation to the time zero sample.* ***b)*** *The expression of hsdM3 in the H. pylori 26695 control mutant at pH 7 or pH 5 at 30 and 60 minutes.* ***c)*** *The expression of hsdM3 in H. pylori ΔarsS at pH 7 or pH 5 at 30 and 60 minutes. Each dot represents a biological replicate, each done in a triplicate. Error bars; standard error of the mean. Statistical analysis via unpaired one-tailed t-test. *; P≤0.05, **; P≤0.01*

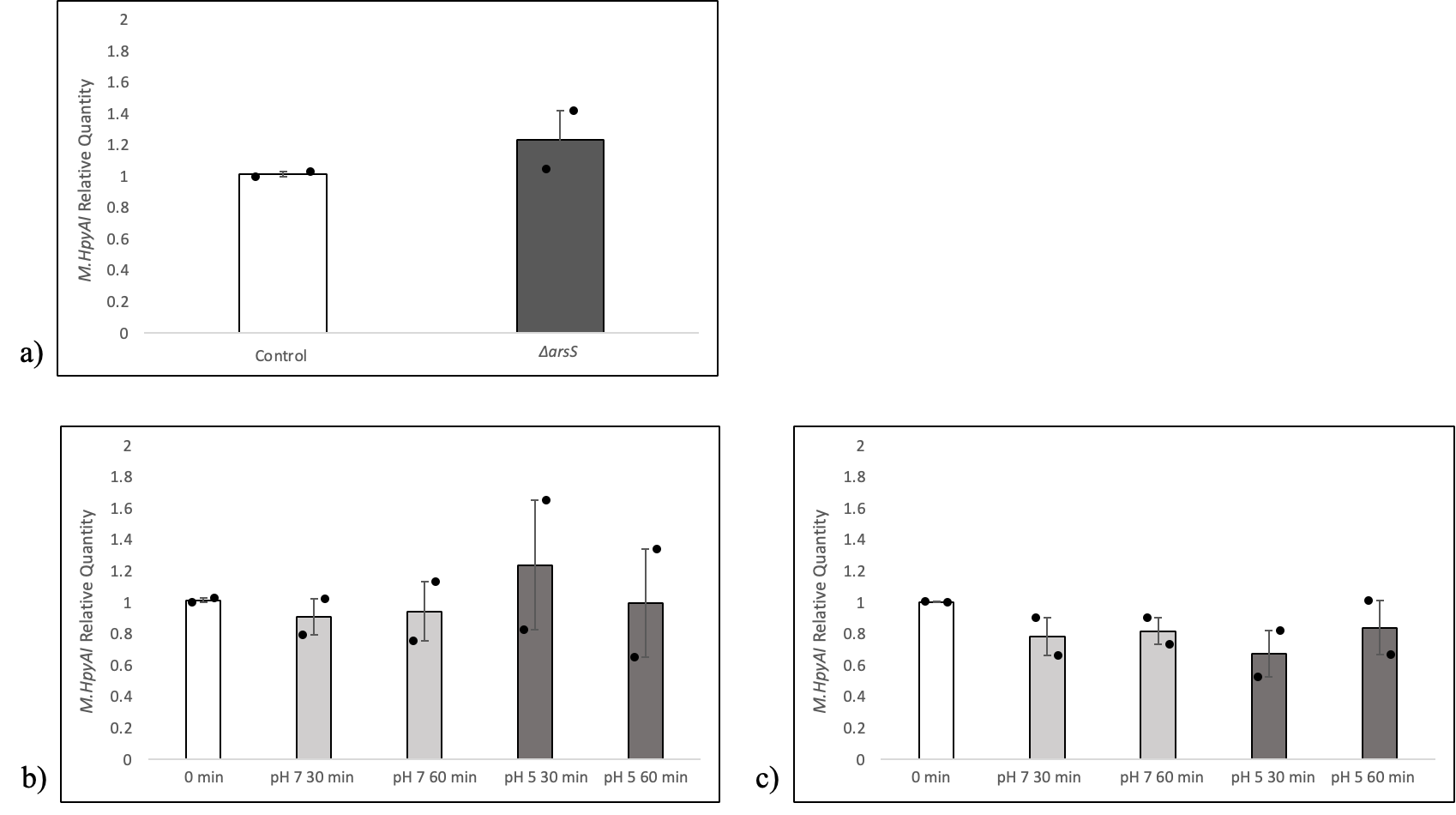

**Supplementary Figure 4.** ***M.HpyAI* (HP1208) transcription is neither acid-sensitive nor ArsS-dependent.** *Both the control H. pylori mutant, possessing an intact arsRS locus, and an isogenic ΔarsS mutant were grown to mid-logarithmic stage of growth and equal aliquots were harvested and resuspended in pH 7 or pH 5 broth.* **a)** *M.HpyAI* *mRNA levels are expressed as relative quantities in relation to the control strain. Both the control H. pylori mutant, possessing an intact arsRS locus, and an isogenic ΔarsS mutant were grown to mid-logarithmic stage of growth and equal aliquots then harvested.* ***b-c)*** *After growth of the control H. pylori mutant and isogenic ΔarsS mutant to mid-logarithmic phase, equal aliquots were harvested and resuspended in pH 7 or pH 5 broth. mRNA levels are expressed as relative quantities in relation to the time zero sample.* ***b)*** *The expression of M.HpyAI in the H. pylori 26695 control mutant at pH 7 or pH 5 at 30 and 60 minutes.* ***c)*** *The expression of M.HpyAI in H. pylori ΔarsS at pH 7 or pH 5 at 30 and 60 minutes. Each dot represents a biological replicate, each done in a triplicate. Error bars; standard error of the mean. Statistical analysis via unpaired one-tailed t-test. *; P≤0.05, **; P≤0.01*

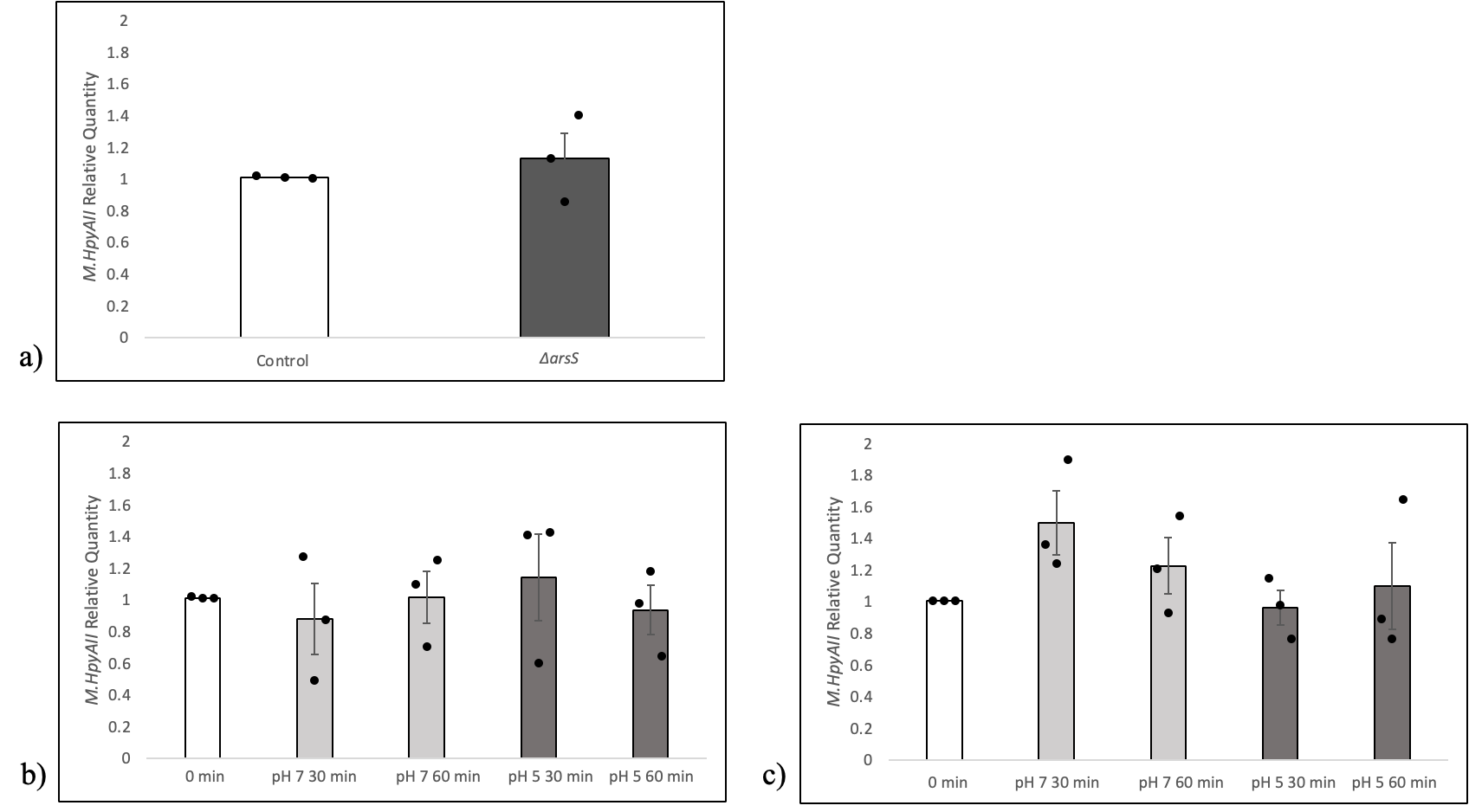

**Supplementary Figure 5.** ***M.HpyAII* (HP1368) transcription is neither acid-sensitive nor ArsS-dependent.** *Both the control H. pylori mutant, possessing an intact arsRS locus, and an isogenic ΔarsS mutant were grown to mid-logarithmic stage of growth and equal aliquots were harvested and resuspended in pH 7 or pH 5 broth.* **a)** *M.HpyAII* *mRNA levels are expressed as relative quantities in relation to the control strain. Both the control H. pylori mutant, possessing an intact arsRS locus, and an isogenic ΔarsS mutant were grown to mid-logarithmic stage of growth and equal aliquots then harvested.* ***b-c)*** *After growth of the control H. pylori mutant and isogenic ΔarsS mutant to mid-logarithmic phase, equal aliquots were harvested and resuspended in pH 7 or pH 5 broth. mRNA levels are expressed as relative quantities in relation to the time zero sample.* ***b)*** *The expression of M.HpyAII in the H. pylori 26695 control mutant at pH 7 or pH 5 at 30 and 60 minutes.* ***c)*** *The expression of M.HpyAII in H. pylori ΔarsS at pH 7 or pH 5 at 30 and 60 minutes. Each dot represents a biological replicate, each done in a triplicate. Error bars; standard error of the mean. Statistical analysis via unpaired one-tailed t-test. *; P≤0.05, **; P≤0.01*

| **Target** | **Forward Probe (5’-3’)** | **Reverse Probe (3’-5’)** | **Reporter (5’-3’)** |
| --- | --- | --- | --- |
| gyrB - DNA Gyrase B subunit (consensus normalizing gene) (*gyrB*) | AAAGCCAGAGAGCTTACAAGGAAAA | CGCCCTCCACTAAAAAGATTTCACT | TTGCCTGGAAAATTAG |
| HP0463- Type I DNA methyltransferase enzyme (*hsdM*1) | ATGAGCCGACTAGAAATGTCAAAATCT | CCTATTTGGTGGGCTAATGCCATT | CCTGTGCCTGCACTTG |
| HP0850 - Type I DNA methyltransferase enzyme (*hsdM*2) | AAAGTGTTAGGCGATAAAAATGTCTCAAAAG | GGCAAAGGTTGTAAGTGGTCAAATT | TCTTGCCCAAAATACC |
| HP1403 - Type I DNA methyltransferase enzyme (*hsdM*3) | CGCGCGCCAGAAAGG | AGCGGTTTTTATTGCCGTCTTTTT | CCTTGCTCGCATCTAT |
| HP1208 - Type II DNA methyltransferase enzyme (*M.HpyAI*) | GCCTTAAAAAAAGCGCTCAAAAAGA | TGCTGATACACTTCGTTAGCGTTTA | AAAGGCGCTGATTTTG |
| HP1368 - Type IIS DNA methyltransferase enzyme (*M.HpyAII*) | ACATGCTAAAAAACAAACCTAAAATGTTCTTACT | GGGTAGCTCCCAAACATCAATCTTT | ACGCGCAAATCCCAC |

**Supplementary Table 2.** *ThermoFisher TaqMan assay sequences used for qRT-PCR probes. The reporters have FAM as the fluorophore at the 5’ end and NQR as the quencher at the 3’ end.*
